# MDM2 case study: Computational protocol utilizing protein flexibility and data mining improves ligand binding mode predictions

**DOI:** 10.1101/054239

**Keywords:** MDM2, autodock, autodock vina, molecular docking, data mining, drug design, molecular dynamics, high throughput virtual screenings

## Abstract

Recovery of the P53 tumor suppressor pathway via small molecule inhibitors of oncoprotein MDM2 highlights the critical role of computational methodologies in targeted cancer therapies. Molecular docking programs in particular, have become essential during computer-aided drug design by providing a quantitative ranking of predicted binding geometries of small ligands to proteins based on binding free energy. In this study, we found improved ligand binding mode predictions of small medicinal compounds to MDM2 based on RMSD values using AutoDock and AutoDock Vina employing protein binding site flexibility. Additional analysis suggests a data mining protocol using linear regression can isolate the particular flexible bonds necessary for future optimum docking results. The implementation of a flexible receptor protocol based on ‘a priori’ knowledge obtained from data mining will improve accuracy and reduce costs of high throughput virtual screenings of potential cancer drugs targeting MDM2.

## 1 Introduction

P53 is a tumor suppressor protein found in the nucleus of cells, which functions to respond to cellular stress by mediating cell-cycle arrest, senescence, or apoptosis in response to DNA damage, oncogene activation, and hypoxia (Zhao et al., 2013)(Vousden and Prives, 2009). Inactivation of the P53 pathway is found in the majority of human cancers and is facilitated by mutation or deletion of the TP53 gene or damage to cellular regulatory mechanisms (Klein and Vassilev, 2004)(Nag et al., 2013). The primary regulator of P53 is murine double minute 2 (MDM2), an E3 ubiquitan ligase protein which binds to P53 marking it for degradation. In damaged cells, over-expression of MDM2 results in reduced levels of P53, initiating the onset of oncogenesis (Wang et al., 2003)(Wang et al., 2001). Chemotherapies attempt to block this interaction and recover tumor suppression activity by introducing small non-peptide molecules designed to target and bind to the P53 binding domain of MDM2 (Wang et al., 2003)(Khoury and Dömling, 2012). Several small molecule inhibitors have been designed from lead compounds discovered using the structure-based virtual screening of chemical libraries aided by docking programs, and many have entered and completed phase 1 cancer drug clinical trials (Bharatham et al., 2014)(Wang et al., 2012). These high throughput virtual screenings (HTVS) evaluate thousands of small molecules and are a cost effective approach designed to rely on fast, accurate predictions, intended to isolate a small number of promising leads as future cancer therapeutics (Grinter and Zou, 2014)(Ellingson et al., 2013).

### 1.1 Molecular docking: A computational methodology aiding drug design

Molecular docking programs represent a critical tool in the early stages of structure-based drug design (SBDD), while providing important insights into molecular binding processes (Meng et al., 2011)(Warner et al., 2012). The focus on MDM2 and subsequent literature has underscored the importance of docking programs such as AutoDock and AutoDock Vina, (henceforth referred to as Vina) for the quick and accurate screening of cancer drug candidates (Houston et al., 2015)(Dhanik et al., 2013). These programs conduct virtual screenings of small molecules from chemical libraries while attempting to manage and resource the vast chemical space of all possible compounds available to be optimized as future cancer drugs (Deligkaris et al., 2014). HTVS can exclude or include available compounds *in silico*, as drug leads based on desired binding geometry often referred to as the binding mode, and binding free energy, a quantitative measure of the binding affinity between molecules (Teodoro et al., 2002). Due to the incredible complexity inherent in simulating the molecular binding process, docking programs introduced a time-independent strategy based on chemical potentials rather than the force calculations associated with classical molecular dynamics (MD), which adopt time dependent Newtonian physics (Plattner and Noé, 2015)(Totrov and Abagyan, 2008). Attempting to strike a balance between computational time and accuracy, docking programs rely on energy evaluations based on assumptions, estimates and empirical knowledge while estimating, rather than calculating, binding free energy (Sousa et al., 2013).

The specific location and binding mode, along with a strong corresponding binding affinity, are the two components of a successful docking (Cosconati et al., 2010). The binding affinity is determined from evaluations based on different molecular interactions and reflects the strength of the preferred non-covalent binding (Meng et al., 2011). Given the complex interactions estimated by the semi-empirical scoring functions often employed by docking programs using simplified free energy models, the docking free energies are generally viewed as un-reliable as a true measure of the binding affinity. This problem arises when the experimental binding energy determined from dissociation constants are quantitatively different from the estimates used for docking experiments (Perola et al., 2004). Nevertheless, docking methodologies in SBDD are widely used as they provide a complimentary technique for the discovery and optimization of lead compounds targeting proteins in addition to DNA intercalates and minor groove binders, designed to disrupt cancerous cell replication (Singh et al., 2016).

### 1.2 Overview of AutoDock and Vina

AutoDock’s efficacy and limitations have been well documented while being shown to provide fast and accurate predictions within 2 Å of the experimentally known binding site for ligands with up to 10 rotatable bonds. However, as the ligands’ rotatable bonds increase, performance decreases, largely due to the exponential increase of possible conformational states (Plewczynski et al., 2011)(Morris et al., 2009). This restriction has been a notable difficulty pertaining to protein-ligand docking and has led to the use of a rigid receptor protocol as standard methodology because of the computational challenges and increased cost posed by incorporating protein flexibility (Abreu et al., 2012)(Spyrakis et al., 2011). This methodology fails to account for side-chain residue movement at binding site interfaces, resulting in a less reliable prediction of the ligands’ docked binding mode (Sotriffer, 2011). We know the accurate, computational simulation of protein conformational changes is critical to improved docking studies because it accounts for changes affecting the final binding geometry (Plattner and Noé, 2015). It has been shown that, when only a rigid receptor conformation is considered, docking studies predict incorrect binding poses for about 50–70% of all ligands (Totrov and Abagyan, 2008). In response, AutoDock introduced a feature providing incorporation of protein flexibility accounting for a small portion of conformational changes upon binding. This additional flexibility supplied to the binding site residues is still subject to the limits imposed by the number of rotatable bonds. Therefore, this feature is limited to the ligand and protein having a total of about 10 rotatable bonds. Although invoking AutoDock’s side-chain flexibility feature accounts for some protein movement, the additional conformational search space associated with a flexible ligand and protein can reduce accuracy and increase computational costs (Antunes et al., 2015).

Vina, a faster and more accurate alternative, was released in 2010. Vina was able to improve accuracy while drastically reducing computation time through effective computer architecture and incorporating a “machine learning” approach for the scoring function (Trott and Olson, 2010). Tested using the same protein-ligand complexes evaluated during the development of AutoDock 4, results show a marked improvement in terms of ligand binding mode accuracy. Vina’s combination of speed and accuracy has made it an ideal program for HTVS and has been used in several research studies and novel docking approaches (Ellingson et al., 2013)(Chang et al., 2010). Although AutoDock 4 and Vina share similarities in the use of an empirically weighted scoring function and global search optimization algorithm, they differ in their local search strategy and scoring function parameters (Trott and Olson, 2010).

### 1.3 Improving ligand binding mode predictions

As aforementioned, the two most significant results from a docking experiment include the binding free energy associated with complex formation and the ligand binding mode prediction. This research is focused on improving the ligand binding mode predictions of AutoDock and Vina through selective flexibility of the ligand and receptor by invoking AutoDock’s feature of protein residue flexibility, while utilizing the speed and accuracy of Vina to reduce computational time. Our re-docking study revealed improved binding mode predictions of small medicinal compounds to MDM2 based on RMSD values from the experimentally known binding site. Analysis of these results was supplemented by a classical MD simulation performed by the Nanoscale Molecular Dynamics program (NAMD) (Phillips et al., 2005). MD simulation programs, such as NAMD, utilize classical Newtonian physics to study the time dependent structure, dynamics, and thermodynamics of biological molecules. The microscopic properties of atomic positions and velocities can be translated into macroscopic quantities including temperature, pressure and volume using statistical mechanics. This enables determination of movement associated with selected binding site residues of the target protein (Adcock and McCammon, 2006). The docking and MD results from this study highlights the importance of modeling protein flexibility for the determination of accurate binding mode predictions of small molecules to MDM2 and may be especially useful for HTVS of potential cancer drugs focusing on different protein and DNA targets. An Additional post-analysis study utilizing linear regression analysis found data mining of docking results may play a crucial role in determining future docking studies by locating the particular rotatable bonds most influencing the binding process. Cancer therapeutic research relies on a critical understanding of bio-molecular interactions enhanced by the effectiveness of computational techniques (Cosconati et al., 2010). Evaluation and optimization, complimented by advanced computer architecture, will aide in reducing the cost of cancer drug development and foster new insights into bio-molecular binding processes.

## 2 Methods and materials

Standard docking experiments employ a rigid receptor/flexible ligand protocol while exploring conformational space within a specified grid box designated by the user. A successful re-docking will be within 2 angstroms of the experimentally known site and correspond to one of the top ranked binding energies (Huang and Zou, 2006)(Bikadi and Hazai, 2009). In short, the sum of the energy of ligand and receptor separately is greater than the total energy when bound together. The difference is the binding free energy. A higher negative energy indicates a deeper potential energy well, a more stable complex, and more likely binding mode (Huey et al., 2007). For this study, only the top ranked binding energy and corresponding RMSD from the known binding site was considered as a data point.

### 2.1 Experimental details

A set of four structures, representing small molecule inhibitors in complex with MDM2 was retrieved from the protein data bank (PDB). PDB codes: 4JRG (12), 3LBK (5), 4IPF (10) and 4ZYI (9). The number of inherent rotatable bonds in each ligand is given in parenthesis. For each complex, our protocol systematically distributed a total of 12 rotatable bonds between the ligand and receptor beginning with 0 flexible bonds for the ligand and 12 for the receptor, and then 1 flexible bond for the ligand and 11 for the receptor and so on, using the notation (0,12) and (1,11) respectively. When the maximum number of rotatable bonds was reached inherent in each ligand, the remainder was transferred to the protein. Docking parameters for all calculations using AutoDock 4.2 were adjusted to 100 runs with 2 × 10^7^ energy evaluations and a grid box size of 60Ǻ 62Ǻ 62Ǻ centered on the ligand with .375Å grid spacing. The grid box for Vina 1.1.2 was set to 27Ǻ 27Ǻ 27Ǻ centered on the ligand with a 1 Å grid spacing and the exhaustiveness was set to 12. All other settings for both programs were kept at default parameters.

All structures were retrieved from the PDB and initially prepared for docking using Chimera software (Pettersen et al., 2004). The ligand was separated from the protein and a short energy minimization was applied to each structure for a duration of 10 steps. Hydrogen atoms were added, water molecules removed, and Gasteiger charges were added to the ligand and protein. The necessary files for docking were prepared in AutoDock tools (ADT). When files are imported into ADT, they are checked for polar hydrogens, water molecules and proper charges. The rotatable bonds of the ligand were altered using the ‘choose torsions’ option. Here, the initially flexible bonds of the ligand can be adjusted for docking and saved. A flexible residue file was also created for the rotatable bonds of the selected protein binding site residues in addition to a separate rigid protein file.

AutoDock results are ranked according to the highest negative binding free energies and corresponding RMSD values from the experimentally determined binding site. Vina presents the binding energies with the top ranked binding free energy always corresponding to a 0 RMSD. The subsequent RMSD values are in relation to this top ranked pose. Determining if Vina and AutoDock can converge on a similar binding mode can be accomplished utilizing visualization software, which can directly compare the experimentally known structure to both programs best prediction. AutoDock and Vina share functional commonalities including the global optimization of the scoring function, pre-calculation of grid maps, and the pre-calculation of distant dependent pair-wise energetics between each atom type. However, they employ a different scoring function and algorithms to obtain binding free energies, and should be considered different programs (Trott and Olson, 2010). Conveniently, both programs utilize the same ligand, receptor, and flexible residue files. These files are included as supplemental materials in addition to the docking parameter files and grid parameter files for AutoDock, and the conf.txt, log, and output files for Vina (S1).

### 2.2 Data mining model

As a guide, we tested a data mining approach using a linear regression model where each of the potential bonds is assigned as one of the parameters to be turned off or turned on (0 or 1). We can then seek the weight factor of each of these parameters by supposing the total cohesive energy as well as RMSD can be quantitatively linked to a linear regression as a superposition of all the parameters:

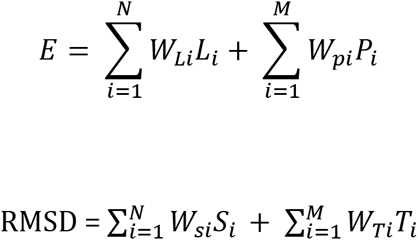

Where:

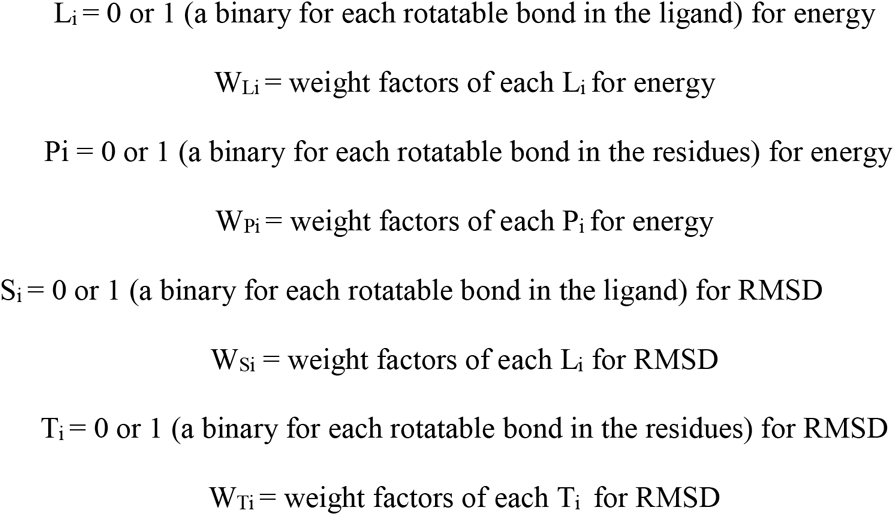

The number of parameters will depend on the total number of rotatable bonds between the ligand and protein. Each parameter can be toggled on or off corresponding to a flexible or rigid bond. This simplified method certainly may not generate fully recoverable linear superposition for a complex docking process; it does provide a guidance based on a given number of allotted rotatable bonds, as to which rotatable bonds that are preferred to be “activated”. The weight functions may also indicate the relative level of importance quantitatively on activating a specific set of rotatable bonds. Further, we can probably identify if there is a potential negative effect in activating certain rotatable bonds toward the total energy and/or RMSD through the formation of negative weight factors. Using this technique, we can locate particular bonds within binding site residues to activate without surveying all possible combinations of protein/ligand flexibility for a particular complex. In general, we can identify the binding site residues of a particular protein, that when made flexible, contribute to optimize the energy and RMSD values associated with a successful docking.

## 3 Results and discussion

The MDM2 protein is a current target for cancer drug development in the form of small molecule inhibitors designed to firmly attach to its P53 binding domain, thus blocking P53/MDM2 interaction. We sought to improve binding mode predictions using the popular docking programs AutoDock and Vina through the selective flexibility of both ligand and binding site residues using four crystallized structures obtained from the PDB. The docking results indicate most accurate binding mode predictions correspond to those configurations supplying maximum protein flexibility. Surprisingly, supplying additional flexibility well past AutoDock’s usual accuracy threshold improves binding site predictions when this additional flexibility is transferred to the protein. The PDB structure 3LBK represents a small molecule in complex with MDM2 that contains only 5 inherent rotatable bonds. AutoDock’s most likely docked pose has it 2.20 Ǻ from the experimentally known site using the standard protocol in contrast to .93 Ǻ, when 9 rotatable bonds are supplied to selected binding site residues and 3 to the ligand (figure 1). A snapshot of each of these two binding modes demonstrates the contrast between AutoDock’s best prediction and the experimentally determined binding geometry (figures 2 and 3). We can see the juxtaposition of predicted and experimental geometries of configuration (5,0) representing the standard protocol and (3,9), representing substantial protein flexibility. The geometry and proximity of the standard protocol docking is not nearly as precise as (3,9), shown by the 2.20 Å RMSD as compared to .93 Å. The ligand bound to MDM2 in structure 4JRG contains 12 rotatable bonds, which is well above AutoDock’s validated limit for a successful docking. Using the standard protocol (12,0), AutoDock’s best prediction is 2.83 Å from the experimentally determined binding mode, while a flexible protein protocol (7,5), yields a prediction within .74 Ǻ (figure 4). Other structures tested, 4IPF and 4ZYI, also show improved binding mode predictions, with lower RMSD values, corresponding to protein flexibility. Complete results from these structures are provided as supplemental materials (S2).

**Figure 1.**
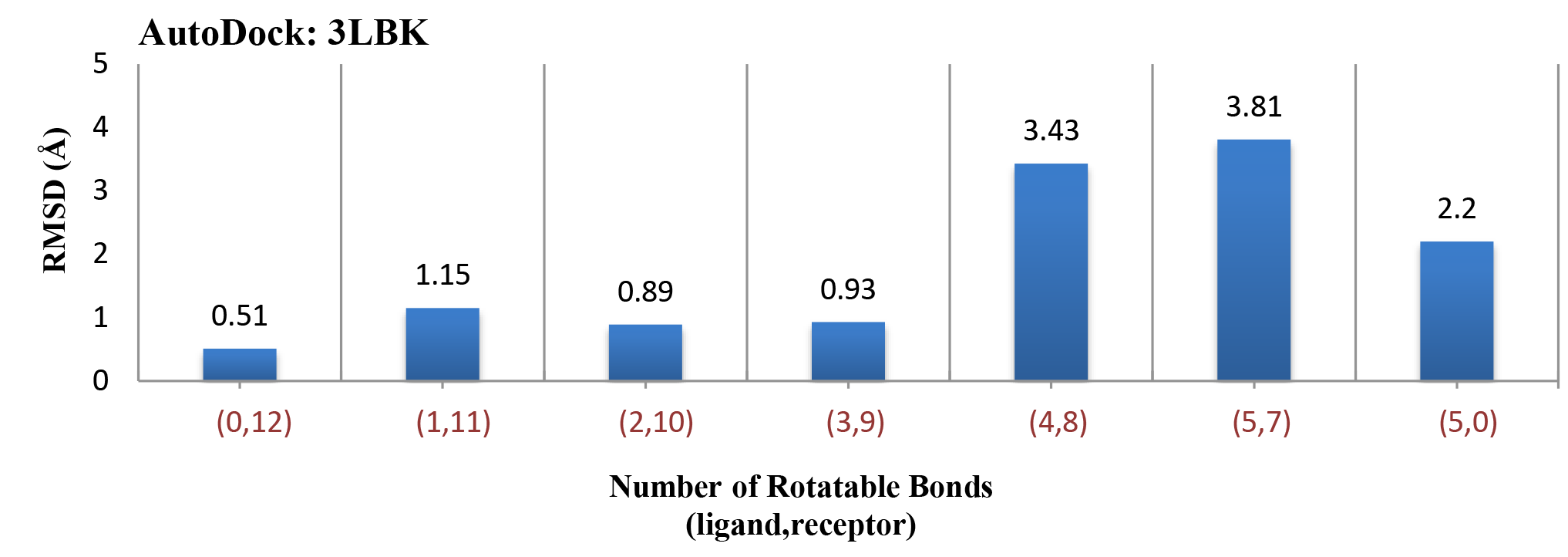
RMSD values corresponding to the top ranked binding energies for all configurations indicate a total rigid ligand (0,12) has the lowest RMSD value of .51 Å. As the number of rotatable bonds become more evenly distributed, binding mode accuracy declines.

**Figure 2.**
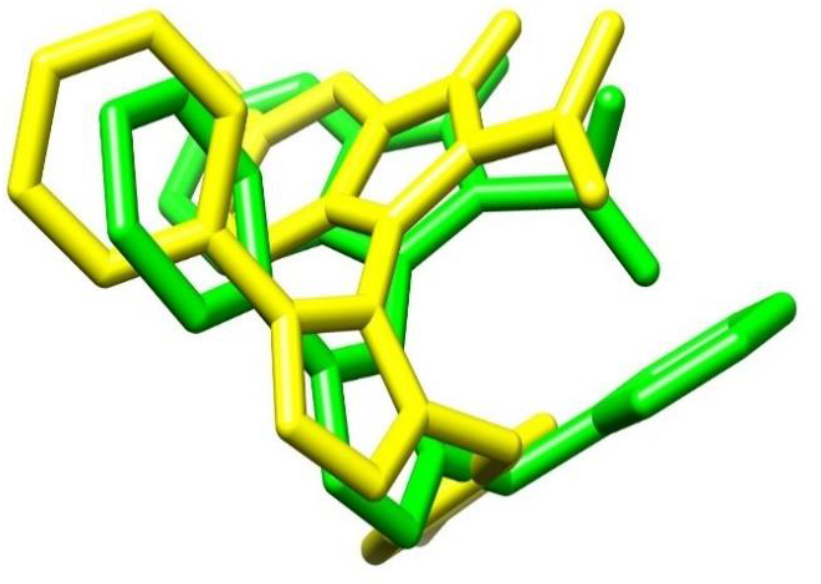
The standard protocol docking pose for 3LBK (5,0) using AutoDock with an RMSD of 2.20 Ǻ. In contrast with the experimentally determined structure.

**Figure 3.**
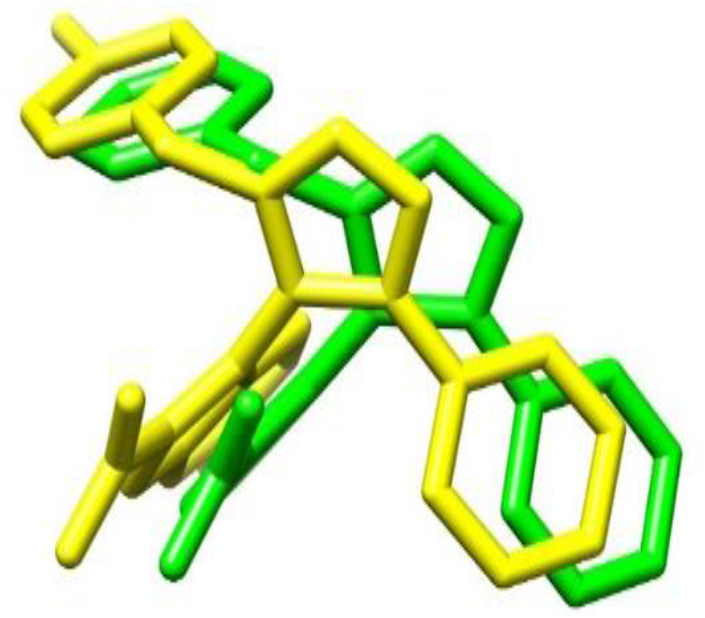
3LBK configuration (3,9) using AutoDock with RMSD of .93 Ǻ. In contrast to the experimentally determined structure.

**Figure 4.**
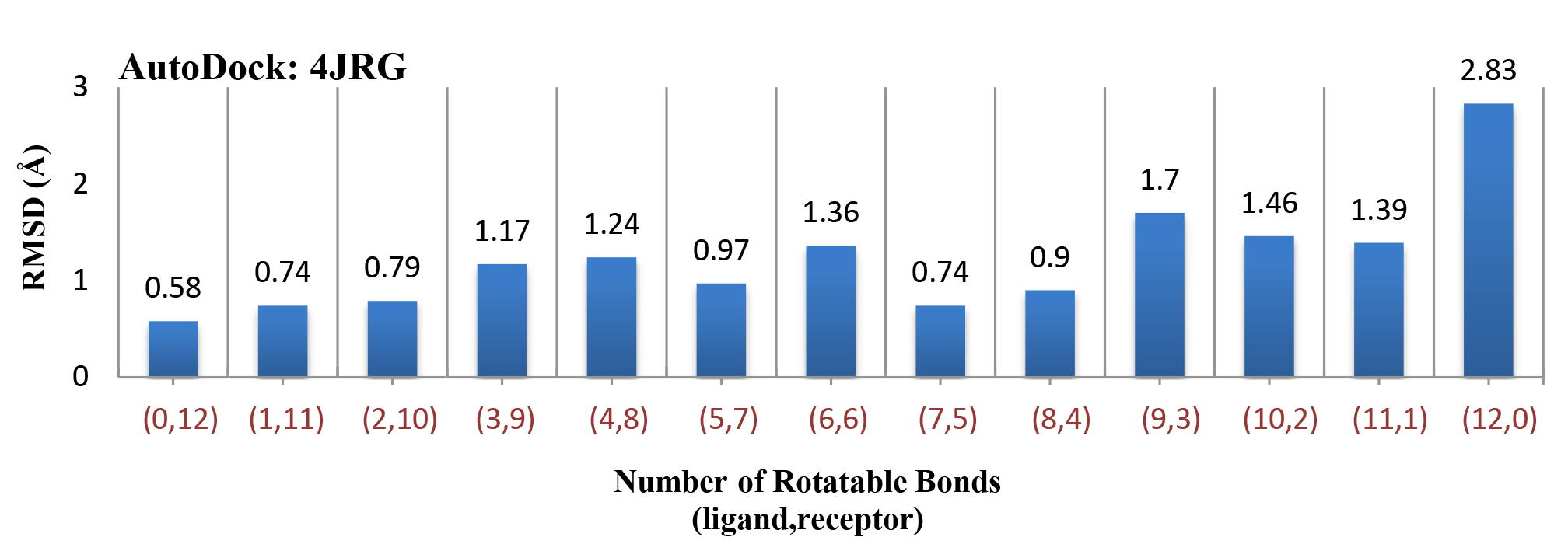
The standard protocol (12,0) configuration shows an RMSD of 2.83 Ǻ with the ligand having 12 rotatable bonds in contrast to a rigid ligand and all 12 rotatable bonds transferred to the MDM2 protein represented by configuration (0,12) with an RMSD of .58 Ǻ.

### 3.1 Vina results

The scoring function and search method developed for Vina has improved speed and binding mode predictions, and is probably better suited for HTVS compared to AutoDock. Vina ranks dockings, but does not associate a RMSD value to the top ranked binding free energy. Hence, Vina results are not charted for this study in terms of RMSD as with AutoDock. Confirmation of a successful docking using Vina and valid comparison to AutoDock must be accomplished using renderings of the binding mode in direct contrast to the experimentally determined structure. Comparing binding free energies is not applicable because AutoDock and Vina use different methods to determine binding free energy with both using many assumptions and estimates. Vina’s most notable advantage is that it mitigates concerns of additional computational time and costs, while allowing for a flexible protein and more accurate binding prediction. Vina’s multithreaded computer architecture can drastically reduce computational time while providing accurate results when docking ligands, as in this study, with 12 rotatable bonds. Vina’s binding prediction for structure 4JRG mirrors AutoDock’s best result of .74 Å (figures 5 and 6). However, Vina’s calculation completed in less than 2 minutes, while the AutoDock calculation lasted 15 hours. Vina, as with AutoDock, also produced a noticeable improvement of binding mode predictions as compared to the standard protocol. Applying a flexible protein protocol when using Vina for HTVS affords the consideration of larger ligands, while providing sufficient accuracy without increased computational costs.

**Figure 5.**
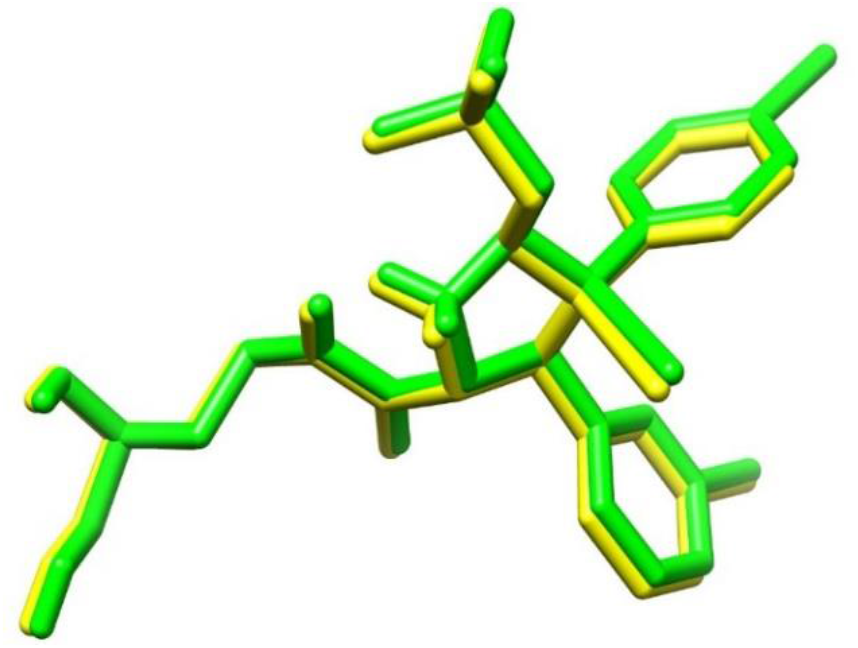
Vina’s prediction of rigid ligand docking of 4JRG (0,12). This binding mode prediction is slightly closer to the experimentally known pose found by AutoDock.

**Figure 6.**
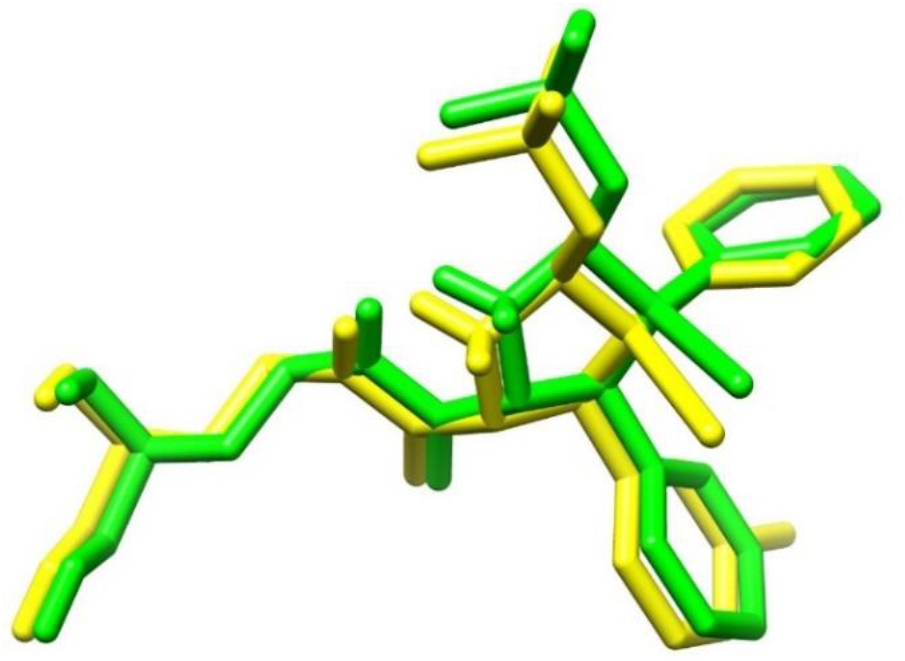
AutoDock’s prediction of rigid ligand docking of 4JRG (0,12). This docked pose is .58 Å from the experimentally determined binding site.

### 3.2 Implications for HTVS

A notable result from this study is the large RMSD values for those configurations representing a completely flexible ligand with limited protein flexibility. For 3LBK, this is docking run (5,7). Although this would seem the ideal distribution of flexibility as it incorporates the inherent flexibility of the ligand and binding protein residues, the RMSD value is 3.81 Å. The same is true of 4JRG, with the (10,2) configuration showing an RMSD of 1.46 Å. This is important because the (5,7) and (10,2) configuration of 3LBK and 4JRG respectively would be applicable to HTVS, as it does not require ligand modification or a change in protein flexibility once the screening starts. This configuration would therefore simulate in part, both protein and ligand binding dynamics. From this study, we find a fully flexible ligand, in combination with selective protein binding site flexibility, fails to optimize the binding mode predictions. Docking results for structures 4IPF and 4ZYI, also show higher RMSD values for this distribution, with values of 1.64 Ǻ and 3.75 Ǻ respectively. A more exhaustive examination of possible rotatable bond combinations simulating ligand and protein flexibility may, for any particular complex, improve the ligand binding mode prediction and improve RMSD values. Due to the large number of possible combinations, this strategy is impractical, and leads to utilizing statistical analysis for assigning protein flexibility for virtual screening studies.

### 3.3 Molecular dynamics analysis

The improved determination of binding geometries while invoking protein flexibility is probably best explained by the physical structure of the MDM2 binding domain which is flanked by residues not embedded within the protein. These residues can fluctuate during the binding process, allowing a ligand to enter while the protein conforms according to interactions determined by the small molecules chemical structure. A Classical MD simulation of the MDM2 protein using the crystallized structure 4IPF was performed by NAMD. A minimization and equilibration simulation allows for the determination of residue mobility before interaction with the ligand from average RMSD values calculated during equilibration. This data provides insight into the protein residue dynamics the ligand encounters as it enters the binding site. The protein was prepared, and necessary files created, using visual molecular dynamics (VMD), the graphical user interface designed to work with NAMD for the preparation, evaluation and visualization of MD simulations. The simulation lasted .5 ns to ensure equilibration with a 1 fs time step at a constant temperature of 310K in an explicit solvent. Results find the residues fluctuate between 1.4 and 3.2 Å, with residues HIS 69, HIS 92 and TYR 96 all moving an average of 3 Å (figure 7). The RMSD values serve to quantify movement of the protein in equilibrium, affirming the importance of modeling residue flexibility during docking experiments. Although the flexibility as modeled by AutoDock and Vina is limited to the rotation of bonds, with bond lengths and angles kept constant, the change observed from flexible binding site residues can be pronounced. The before and after snapshot of selected residues indicates the movement necessary to produce a successful docking within .49 Å of the known structure (figure 8). This suggests standard protocol docking employing ligand flexibility may not be as essential to predicting an accurate binding mode as modeling the protein conformational changes accommodating a small molecule during the binding process. A short MD trajectory movie of 4IPF simulating protein movement is provided in the supplementary material (S3).

**Figure 7.**
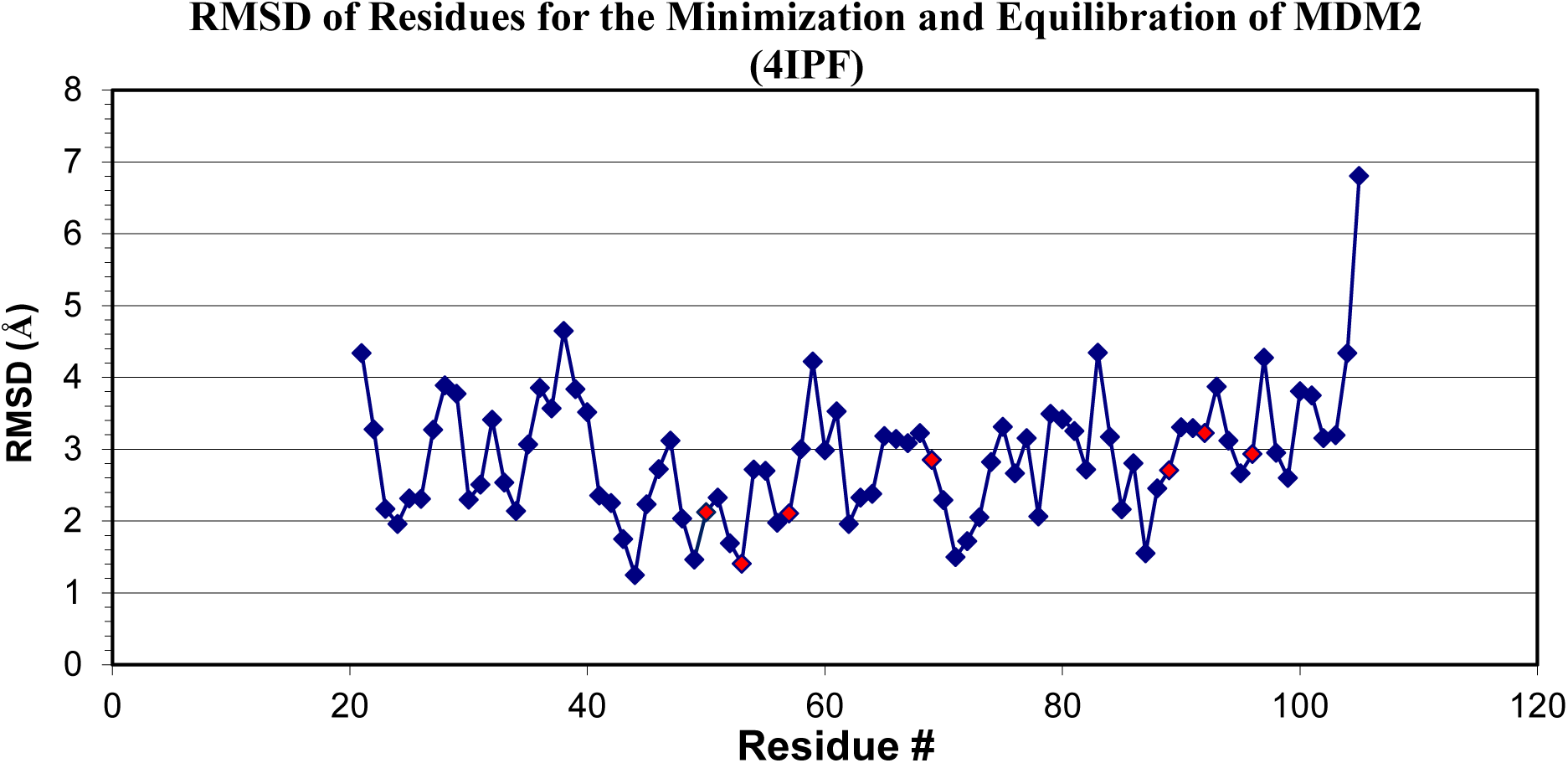
The MDM2 protein as represented by crystal structure 4IPF was selected to evaluate fluctuation of residues providing quantitative data for a better understanding of the binding site dynamics. Markers indicate flexible binding site residues.

**Figure 8.**
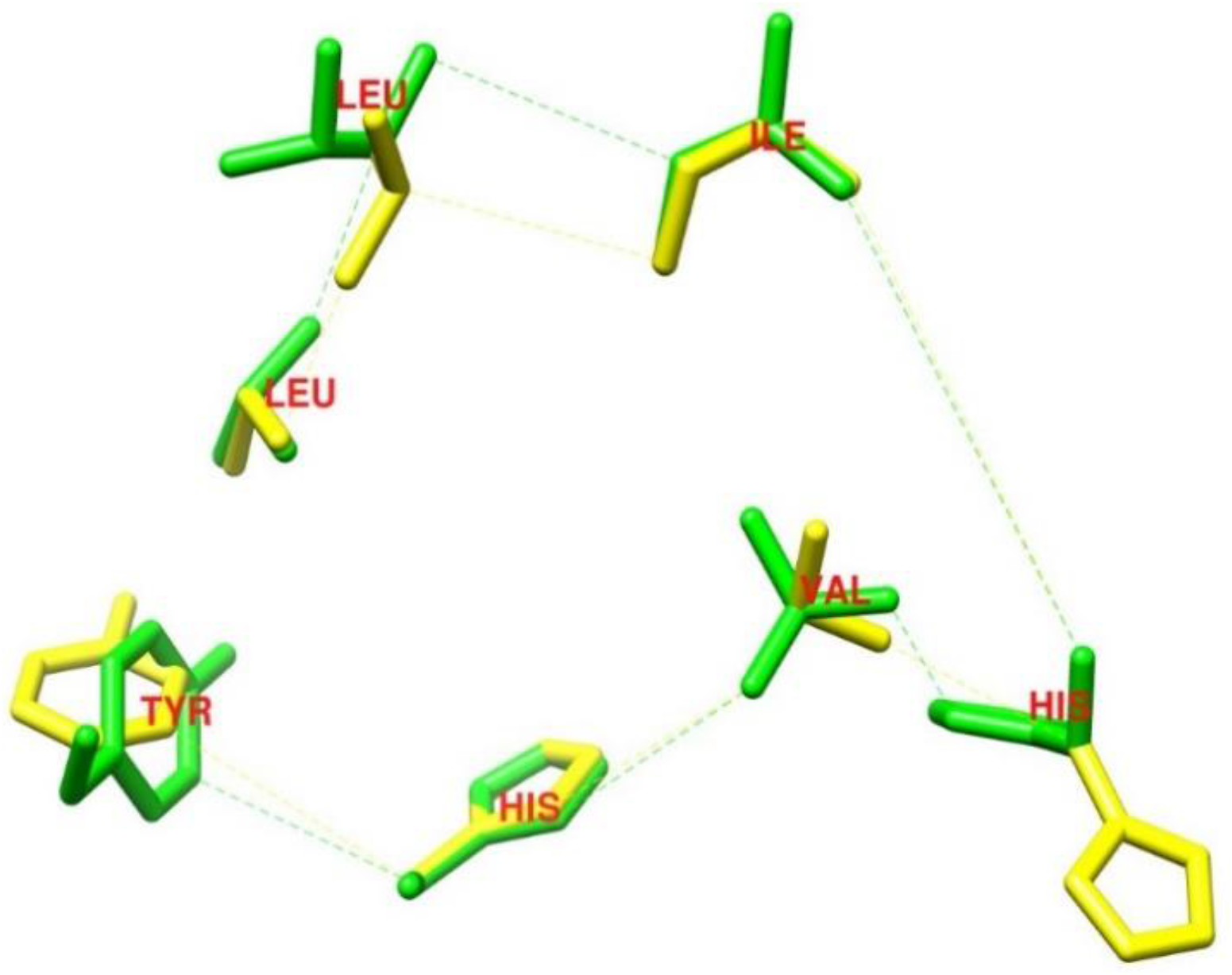
Final positions of flexible binding site residues for 4IPF (0,12) in contrast to pre-docking positions. Residues LEU, TYR, VAL and HIS have shifted considerable from the experimentally determined geometry. Due to the time independent nature of docking, we can only capture before and after states of the residues. The protein and ligand have been removed for clarity.

### 3.4 Data mining study

Docking results for structure 3LBK were used as a data set for a linear regression analysis to determine the optimal flexibility distribution for the protein and ligand (table 1). Ideally, we want to isolate the critical rotatable bonds responsible for maximizing the magnitude of the total energy and/or minimizing the RMSD. In a way, it is a “genomic” approach, to ID the rotatable bonds. For 3LBK, overall we have identified P2, P4, P8 and P9 as the rotatable bonds that can be used to construct a number of simple linear models describing the values of RMSD and Total Energy (see the linear formulas below and corresponding figure 9).

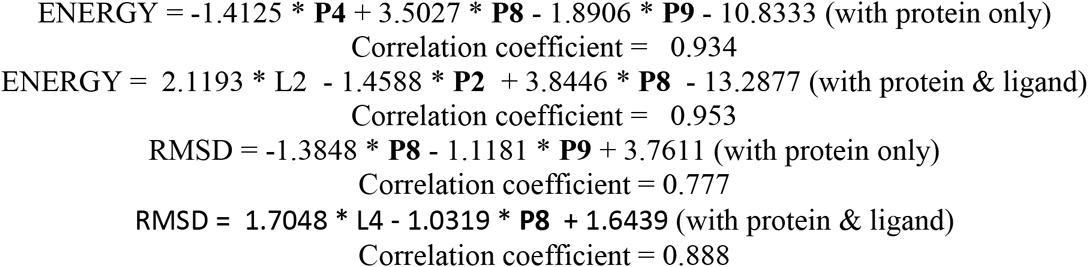

**Figure 9.**
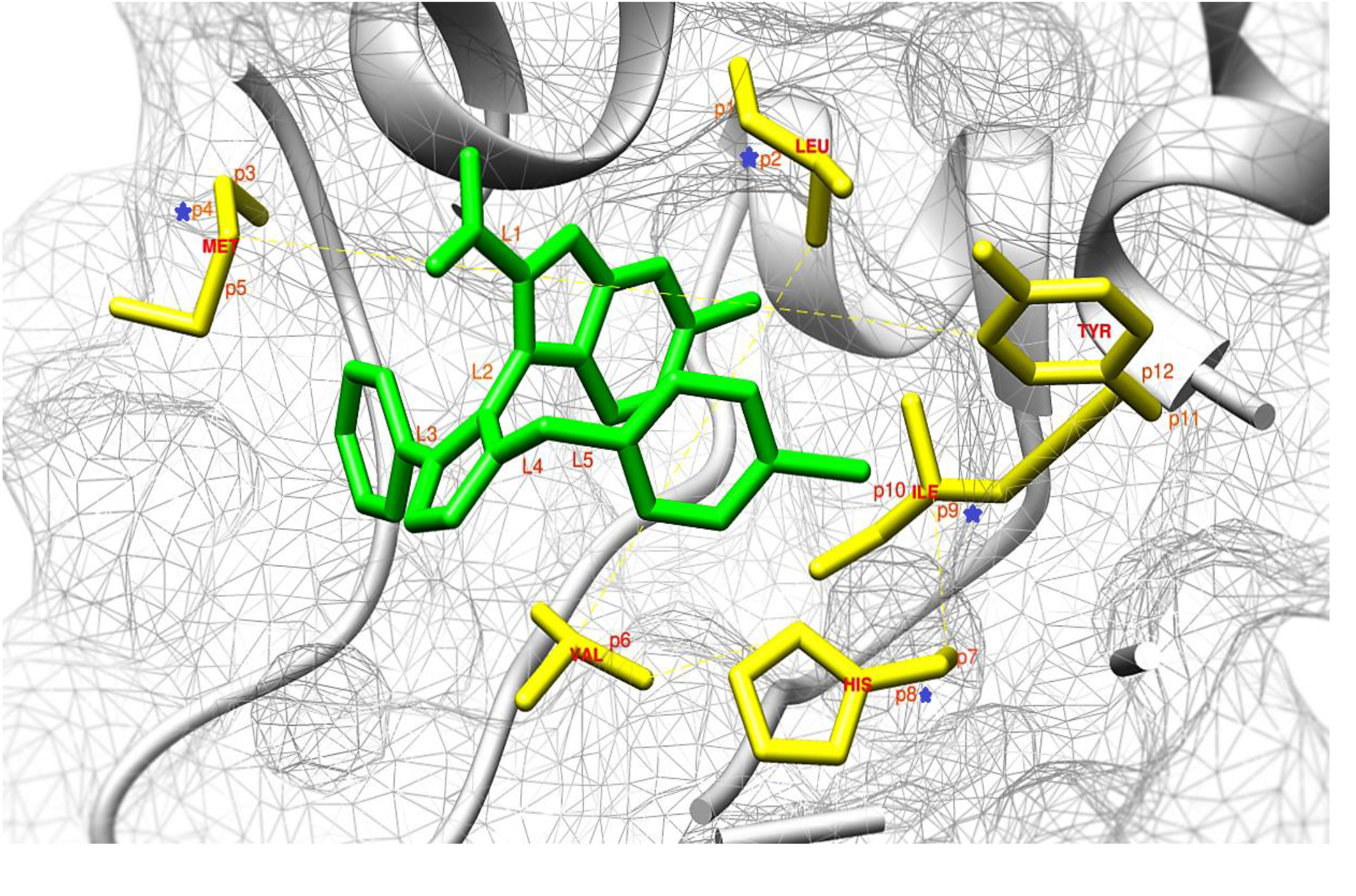
Structure 3lbk representing a small molecule inhibitor in complex with MDM2. Linear regression results from 12 docking studies found particular rotatable bonds help to improve energy and RMSD values. Rotatable bonds from binding site residues are labeled P1 through P12. Blue stars denote those bonds weighted favorably for flexibility.

**Table 1.**
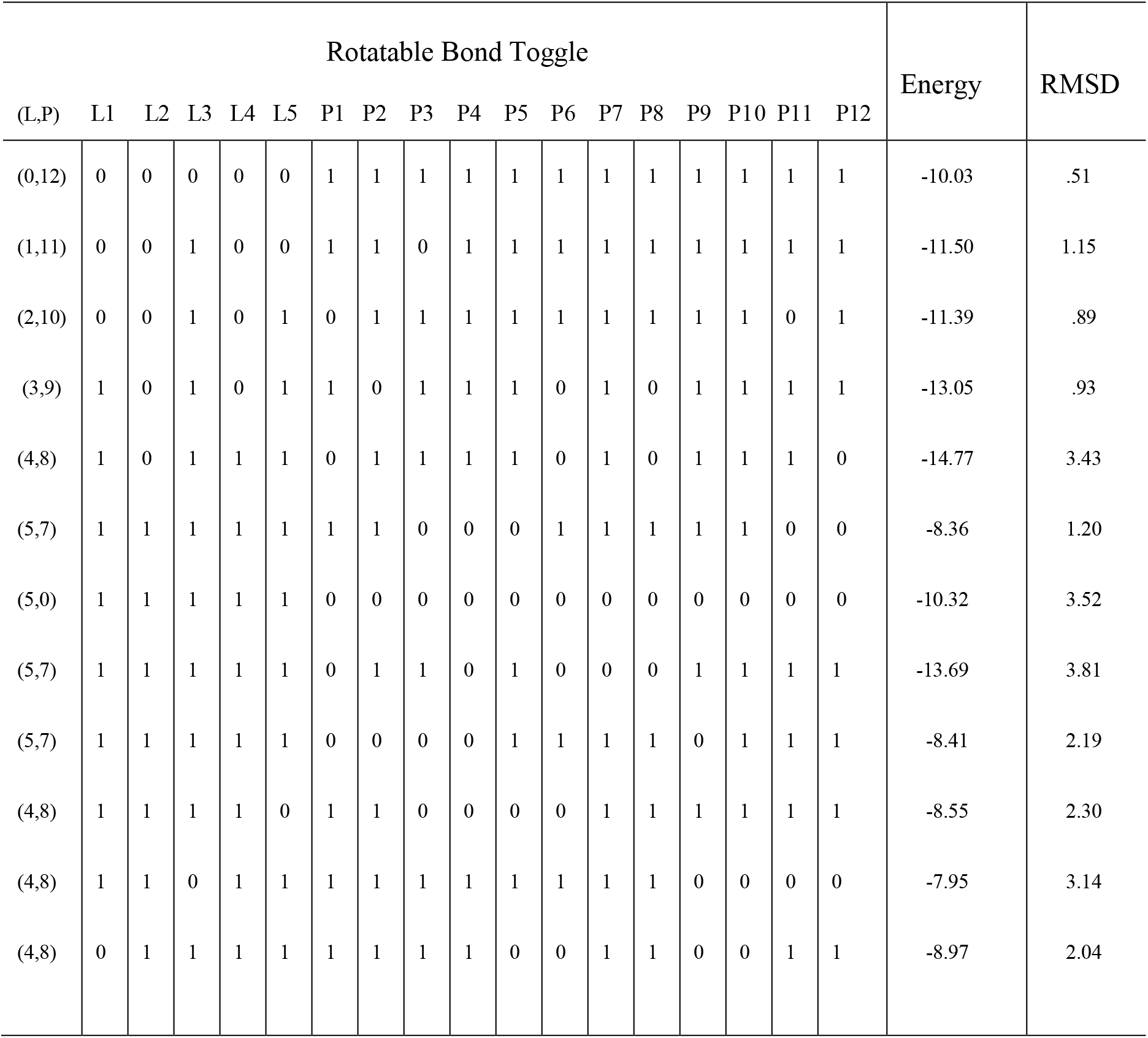
Structure 3lbk bond toggle displays the activation status of each bond for each configuration in addition to the correspond energy and RMSD values.

This finding would allow us to assign these four rotatable bonds to be preferable active in future docking procedures. A close examination of these four rotatable bonds show that these are not necessarily the bonds that govern the terminating ends of their residues. For example, as shown in Figure 9, the P4 affects the connection between CB-CG in MET residue, positioned in the middle portion of the residue. Similarly, P9 which connect CA and BC in ILE residue. This suggests that it is critical to add and optimize the flexibility in this protein.

## 4 Conclusion

Improved binding mode predictions of small molecule inhibitors targeting MDM2 was achieved using AutoDock and Vina employing a systematic distribution of 12 rotatable bonds between the ligand and protein. This study found activating bonds of selected binding site residues produced lower RMSD values when compared to standard rigid receptor docking. Additional analysis discovered considerable movement of key residues selected for docking illustrated by a MD simulation. Further, the data mining method we adopted shows promise as a cost effective method for facilitating ‘a priori’ knowledge necessary for the optimized selection of flexible bonds. Adopting a flexible protein docking protocol aided by statistical analysis for future medicinal studies of MDM2 using Vina in particular, will enable accurate, fast predictions of small molecule inhibitors of MDM2. Future studies will determine if this protocol may be applied to additional protein targets of medicinal interest.

## References

Abreu, R.M.V, Froufe, H.J.C., Queiroz, M.-J.R.P., Ferreira, I.C.F.R., 2012. Selective flexibility of side-chain residues improves VEGFR-2 docking score using AutoDock Vina. Chem. Biol. Drug Des. 79, 530–4.

Adcock, S.A., McCammon, J.A., 2006. Molecular dynamics: survey of methods for simulating the activity of proteins. Chem. Rev. 106, 1589–615.

Antunes, D.A., Devaurs, D., Kavraki, L.E., 2015. Understanding the Challenges of Protein Flexibility in Drug Design. Expert Opin. Drug Discov. 10, 1–17.

Bharatham, N., Bharatham, K., Shelat, A.A., Bashford, D., 2014. Ligand binding mode prediction by docking: mdm2/mdmx inhibitors as a case study. J. Chem. Inf. Model. 54, 648–59.

Bikadi, Z., Hazai, E., 2009. Application of the PM6 semi-empirical method to modeling proteins enhances docking accuracy of AutoDock. J. Cheminform. 1, 15.

Chang, M.W., Ayeni, C., Breuer, S., Torbett, B.E., 2010. Virtual Screening for HIV Protease Inhibitors: A Comparison of AutoDock 4 and Vina. PLoS One 5, e11955.

Cosconati, S., Forli, S., Perryman, A.L., Harris, R., Goodsell, D.S., Olson, A.J., 2010. Virtual Screening with AutoDock: Theory and Practice. Expert Opin. Drug Discov. 5, 597–607.

Deligkaris, C., Ascone, A.T., Sweeney, K.J., Greene, A.J.Q., 2014. Validation of a computational docking methodology to identify the non-covalent binding site of ligands to DNA. Mol. Biosyst. 10, 2106–25.

Dhanik, A., McMurray, J.S., Kavraki, L.E., 2013. DINC: A new AutoDock-based protocol for docking large ligands. BMC Struct. Biol. 13, S11.

Ellingson, S.R., Smith, J.C., Baudry, J., 2013. VinaMPI: facilitating multiple receptor high-throughput virtual docking on high-performance computers. J. Comput. Chem. 34, 2212–21.

Grinter, S.Z., Zou, X., 2014. Challenges, applications, and recent advances of protein-ligand docking in structure-based drug design. Molecules 19, 10150–76.

Houston, D.R., Yen, L.-H., Pettit, S., Walkinshaw, M.D., 2015. Structure- and Ligand-Based Virtual Screening Identifies New Scaffolds for Inhibitors of the Oncoprotein MDM2. PLoS One 10, 1–19.

Huang, S.-Y., Zou, X., 2006. Efficient molecular docking of NMR structures: Application to HIV-1 protease. Protein Sci. 16, 43–51.

Huey, R., Morris, G.M., Olson, A.J., Goodsell, D.S., 2007. A semiempirical free energy force field with charge-based desolvation. J. Comput. Chem. 28, 1145–52.

Khoury, K., Dömling, A., 2012. P53 mdm2 inhibitors. Curr. Pharm. Des. 18, 4668–78.

Klein, C., Vassilev, L.T., 2004. Targeting the p53-MDM2 interaction to treat cancer. Br. J. Cancer 91, 1415–9.

Meng, X.-Y., Zhang, H.-X., Mezei, M., Cui, M., 2011. Molecular docking: a powerful approach for structure-based drug discovery. Curr. Comput. Aided. Drug Des. 7, 146–57.

Morris, G.M., Goodsell, D.S., Pique, M.E., Lindstrom, W. “Lindy,” Huey, R., Stefano, Forli, 2009. AutoDock 4.2 User Guide.

Nag, S., Qin, J., Srivenugopal, K.S., Wang, M., Zhang, R., 2013. The MDM2-p53 pathway revisited. J. Biomed. Res. 27, 254–71.

Perola, E., Walters, W.P., Charifson, P.S., 2004. A Detailed Comparison of Current Docking and Scoring Methods on Systems of Pharmaceutical Relevance. PROTEINS Struct. Funct. Bioinforma. 56, 235–249.

Pettersen, E.F., Goddard, T.D., Huang, C.C., Couch, G.S., Greenblatt, D.M., Meng, E.C., Ferrin, T.E., 2004. UCSF Chimera--a visualization system for exploratory research and analysis. J. Comput. Chem. 25, 1605–12.

Phillips, J.C., Braun, R., Wang, W., Gumbart, J., Tajkhorshid, E., Villa, E., Chipot, C., Skeel, R.D., Kalé, L., Schulten, K., 2005. Scalable molecular dynamics with NAMD. J. Comput. Chem. 26, 1781–802.

Plattner, N., Noé, F., 2015. Protein conformational plasticity and complex ligand-binding kinetics explored by atomistic simulations and Markov models. Nat. Commun. 6, 7653.

Plewczynski, D., Lazniewski, M., Augustyniak, R., Ginalski, K., 2011. Can we trust docking results? Evaluation of seven commonly used programs on PDBbind database. J. Comput. Chem. 32, 742–55.

Singh, S., Das, T., Awasthi, M., Pandey, V.P., Pandey, B., Dwivedi, U.N., 2016. DNA topoisomerase-directed anticancerous alkaloids: ADMET-based screening, molecular docking, and dynamics simulation. Biotechnol. Appl. Biochem. 63, 125–37.

Sotriffer, C.A., 2011. Accounting for induced-fit effects in docking: what is possible and what is not? Curr. Top. Med. Chem. 11, 179–91.

Sousa, S.F., Ribeiro, A.J.M., Coimbra, J.T.S., Neves, R.P.P., Martins, S.A., Moorthy, N.S.H.N., Fernandes, P.A., Ramos, M.J., 2013. Protein-Ligand Docking in the New Millennium - A Retrospective of 10 Years in the Field. Curr. Med. Chem. 20, 2296–2234.

Spyrakis, F., BidonChanal, A., Barril, X., Luque, F.J., 2011. Protein flexibility and ligand recognition: challenges for molecular modeling. Curr. Top. Med. Chem. 11, 192–210.

Teodoro, M.L., Phillips, G.N., Kavraki, L.E., 2002. Molecular docking: a problem with thousands of degrees of freedom, in: Proceedings 2001 ICRA. IEEE International Conference on Robotics and Automation (Cat. No.01CH37164). IEEE, pp. 960–965.

Totrov, M., Abagyan, R., 2008. Flexible ligand docking to multiple receptor conformations: a practical alternative. Curr. Opin. Struct. Biol. 18, 178–84.

Trott, O., Olson, A.J., 2010. AutoDock Vina: improving the speed and accuracy of docking with a new scoring function, efficient optimization, and multithreading. J. Comput. Chem. 31, 455–61.

Vousden, K.H., Prives, C., 2009. Blinded by the Light: The Growing Complexity of p53. Cell 137, 413–31.

Wang, H., Nan, L., Yu, D., Agrawal, S., Zhang, R., 2001. Antisense anti-MDM2 oligonucleotides as a novel therapeutic approach to human breast cancer: in vitro and in vivo activities and mechanisms. Clin. Cancer Res. 7, 3613–24.

Wang, H., Yu, D., Agrawal, S., Zhang, R., 2003. Experimental therapy of human prostate cancer by inhibiting MDM2 expression with novel mixed-backbone antisense oligonucleotides: In vitro and in vivo activities and mechanisms. Prostate 54, 194–205.

Wang, S., Zhao, Y., Bernard, D., Aguilar, A., 2012. Protein-Protein Interactions. Top Med Chem 8, 57–80.

Warner, W.A., Sanchez, R., Dawoodian, A., Li, E., Momand, J., 2012. Identification of FDA-approved Drugs that Computationally Bind to MDM2. Chem. Biol. Drug Des. 80, 631–637.

Zhao, Y., Bernard, D., Wang, S., 2013. Small Molecule Inhibitors of MDM2-p53 and MDMX-p53 Interactions as New Cancer Therapeutics. Biodiscovery 4, 1–15.

